# Cycling reduces the entropy of neuronal activity in the human adult cortex

**DOI:** 10.1101/2024.01.31.578253

**Authors:** Iara Beatriz Silva Ferré, Gilberto Corso, Gustavo Zampier dos Santos Lima, Sergio Roberto Lopes, Mario André Leocadio-Miguel, Lucas G S França, Thiago de Lima Prado, John Araújo

## Abstract

Electroencephalogram (EEG) data is often analyzed from a Brain Complexity (BC) perspective, having successfully been applied to study the brain in both health and disease. In this study, we employed recurrence entropy to quantify BC associated with the neurophysiology of movement by comparing BC in both resting state and cycling movement. We measured EEG in 24 healthy adults, and placed the electrodes on occipital, parietal, temporal and frontal sites, on both the right and left sides. EEG measurements were performed for cycling and resting states and for eyes closed and open. We then computed recurrence entropy for the acquired EEG series. Our results show that open eyes show larger entropy compared to closed eyes; the entropy is also larger for resting state, compared to cycling state for all analyzed brain regions. The decrease in neuronal complexity measured by the recurrence entropy could explain the neural mechanisms involved in how the cycling movements suppress the freezing of gate in patients with Parkinson’s disease due to the constant sensory feedback caused by cycling that is associated with entropy reduction.

## Introduction

Richard Caton once said that the brain speaks for itself. This statement led the pioneer to unveil the waves of electrical potentials in the brain that later inspired Hans Berger in the development of the electroencephalogram (EEG) as a means of studying the human brain at rest in the early 1930s [1, 2]. It was only in 1949, though, that the role of simple voluntary movements in reducing the power of alpha and beta rhythms in sensorimotor regions was described [3]. In turn, due to difficulties in quantifying brain activity during complex movements, the same phenomenon was only reported recently for human gait [4, 5], as measuring brain potentials during movement is not a simple task.

Walking involves important cognitive processes governing neural networks incorporating sensory information, motor planning and execution [6]. Whilst an essential component of human life, this complexity is the reason our walking capacity is greatly sensitive to ageing [7] and can be severely compromised in neurological disorders, such as Parkinson’s disease [8] - which, amongst other symptoms, increases the risk of falls [9]. Compared to walking, cycling comprises a cyclical and repetitive movement, consisting of the complete rotation of the pedal axis around the central axis of the bicycle. While the motor planning and execution of cycling might have similarities to the cortical control of walking, it is a less cognitively demanding movement. For instance, the need for trunk balance is significantly reduced, when compared to walking [4]. This discrepancy served as the motivation for the investigation of the EEG signal between cycling and walking. Storzer et al. [10] contrasted the EEG dynamics involved in cycling and walking in healthy volunteers. Compared to walking, cycling resulted in a reduction of neuronal recruitment for the execution of the movement and therefore corresponds to a simpler dynamics than walking.

Our motivation for performing this study is to understand how cycling interferes with the phenomenon of gait, which could provide tokens on the mechanisms of freezing in Parkinson’s Disease (PD) [11]. The degeneration in the substantia nigra, the hallmark of PD, can lead to difficulties in walking or even freezing of gait [12], a clinical phenomenon characterized by brief episodes of inability to take a step associated with short steps that normally occur at the beginning of gait or when turning during walking, greatly impairing the mobility of patients, and resulting in the increased risk of falls and drastically reducing their quality of life [9]. Recently, researchers have found that the simple movement of cycling decreases PD motor signals while patients ride a bicycle [13]. Also, dynamic cycling increases sensory input to the motor control of movement in Parkinson’s disease patients, which may be related to improvements in motor speed and quality [14].

In this study, we investigated Brain complexity (BC) and its association with the neurophysiological context during the cycling in healthy volunteers. To implement this task, we contrasted the scalp electroencephalogram (EEG) of individuals while they were cycling and during resting state. Furthermore, we analyzed the EEG through a BC measure based on entropy. This concept of BC was introduced to analyse global aspects of brain activity [15]. Some results about BC are well established, for instance, during slow wave sleep or when the eyes are closed during wakefulness the BC decreases [16, 17]. Entropy has been proposed as a tool to distinguish the EEG of Parkinson’s disorder patients from controls [18], as well as to identify the transitions from walking to freezing in such patients [19].

Therefore, we evaluated brain dynamics during cycling, which results in changes in BC when compared to walking. Here we hypothesise that cycling itself does reduce BC when compared to rest. We test this hypothesis with the aid of an entropy quantifier, estimated from the EEG of resting versus cycling in different brain areas, as well as during open and closed eyes, taking brain lateralisation into consideration. Finally, we discussed the findings of the paper in the context of movement neurophysiology and potential mechanisms impaired in Parkinson’s disease.

## Materials and methods

### Experimental Setup

#### Sample

The sample consisted of 24 adult volunteers, 13 women (54%; mean age = 21.30; SE = 0.49) and 11 men (46%; mean age = 21.63; SE = 0.87). We performed an assessment of the subjects through an interview about the clinical conditions and measurements of blood pressure and heart rate before the experiment. Subjects who reported any cardio-respiratory or neurological disease or who had blood pressure values above 90 mmHg and 140 mmHg, for diastolic and systolic blood pressure, respectively, were excluded from the sample.

#### Legal and ethical aspects

The present study was analysed by the Research Ethics Committee (CEP) of Universidade Federal do Rio Grande do Norte, and approved under CAAE 02979318.0.0000.5537. In order to participate in our project, the volunteers expressed their consent to participate in the research by signing the informed consent form (written consent) according to the Helsinki Declaration. The study recruited participants from March 1st to June 14th, 2019.

#### Electroencephalographic evaluation

For the acquisition of cortical electrical activity data, we used the electroencephalography technique, which is a recording of the electrical brain activity with electrodes fixed to the individual’s scalp. For each individual, the duration of the recordings was 2 minutes for resting and 2 minutes for cycling states. This is a non-invasive technique and can be applied repeatedly. For signal acquisition, Ag-AgCl electrodes were positioned on the scalp according to the 10-20 system. An electroencephalographic assembly was made composed of eight electrodes in bimodal assembly, the following pairs being F3-Fz, F4-Fz, C3-Cz, C4-Cz, P3-Pz, P4-Pz, O1-A2 and O2-A1. For the sake of understanding, we named the electrodes as follow: F, C, P and O, in addition to left and right sides. For the placement of the electrodes, the subjects only had to stop using hair creams on the day of the experiment to facilitate fixation of the EEG cap on the scalp and data collection. An abrasive paste for skin asepsis was used in the fixation sites of each electrode, and all electrodes were affixed with a conductive paste at each pre-established site of the scalp.

Data were collected at a sampling rate of 1000 Hz using a PowerLab 8/30 system (AdInstruments, Australia). ECG data were recorded on a PowerLab 26T system. Both systems were integrated, and the data was recorded using Labchart 7 Pro Software (AdInstruments, Australia). Data were captured with a band pass filter from 1 to 100Hz. For analysis, a band-stop filter of 59-61 Hz was applied to remove noise from the electrical network and, during cycling activity, a band-pass filter of 3 to 35 Hz was applied.

#### The bicycle model

There are three main types of stationary bikes: horizontal, upright and spinning. In this experiment, we used a horizontal bicycle, which is generally used as a form of aerobic exercise for cardiac rehabilitation, weight loss, and as a form of stress testing. The horizontal bike reduces possible sensory interference and the risk of impacts for individuals who participate in the activity, such as the risk of falls. Moreover, the individuals cycling in the horizontal bike are more stable and as a consequence the skin electrodes have a better connection. In fact, the electrophysiology artifacts related to a weak skin electrode connection have a great impact in the quality of the measurements. The bicycle model used in this project is the MAX-H by Dream Fitness (Brazil) (Supplementary Figure 1).

### Data

We excluded individuals which showed artefacts in the recorded EEG, from the initial 24 individuals, 6 individuals were excluded. The excluded individuals showed erratic jumps in the electric signal that are typical artefacts that result from a poor skin-electrode contact.

From each individual, a total of 32 electrophysiologic recordings were obtained. These recordings correspond to eight electrode signals measured in four distinct behavioral states. The eight electrodes correspond to the places O, P, F and C for right and left sides. Moreover, each of the individuals was registered at rest and cycling, as well as with open and closed eyes.

### Complexity analysis

Out of several complexity indices used to measure complexity of time series in previous research, e.g., entropy, Lyapunov exponent, or fractal dimension [20–23]; entropy is a long-standing tool to explore complex phenomena [24]. Information entropy is defined according to the following relation in equation 1.

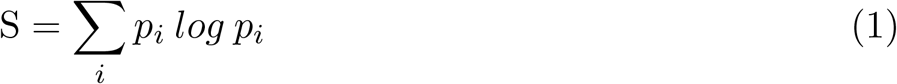

for *p*_*i*_ a probability associated with the time series. In the original work of Shannon [25], the probability is associated with information carried by the time series. In neuroscience literature, the probability is usually related to the amplitude of the spectral representation of the signal [26]. In our work, we skipped the spectral analysis and computed entropy using the recurrence method [27, 28]. This novel technique has been successfully employed to capture signal complexity by evaluating the periodicity of a time series. Recurrence analysis has been widely employed in dynamical systems research and time series analyses [29] as it requires a smaller data amounts to return reliable results [30, 31]. In our case, we split each electrophysiologic recording into 100 pieces to compute the recurrence indices.

## Statistics

To assess statistical significance of our results, we fit a linear mixed effects models, according to GLM1: *MedianEntropy* ∼*Eyes*(*Open/Closed*) + (1|*Subject*). This first analysis was performed as a validation step and compared with previous results reported elsewhere. We then fit a second linear model considering the cycling effect according to GLMM2:

*MedianEntropy* ∼ *State*(*Cycling/Rest*) * *Eyes*(*Open/Closed*) + (1 | *Subject*). In both cases, the statistical significance of the variables of interest was evaluated with two-sided 10,000 repetitions permutation tests.

## Data and code availability

Data acquired in the context of this study, the code used to generated the results, figures and statistics in this article are available in the Open Science Framework repository at https://doi.org/10.17605/OSF.IO/CW87P and at https://github.com/AlgosL/cycling-entropy.git. The script for entropy estimation is available at https://github.com/AlgosL/maxEntropy.

## Results

### Reduced cortical entropy with eyes closed

To validate the recurrence entropy methodology, we compared the signal complexity in a region of the brain where we expect a strong signal response. The entropy of the electrode signal from the right side of the occipital region is shown in Fig. 1. In this situation, all individuals are at rest, and we just compared eyes open and closed. As expected, the open eye shows larger entropy compared to closed eyes *t*(16) = ™2.657; *p* = 0.015.

**Fig 1.**
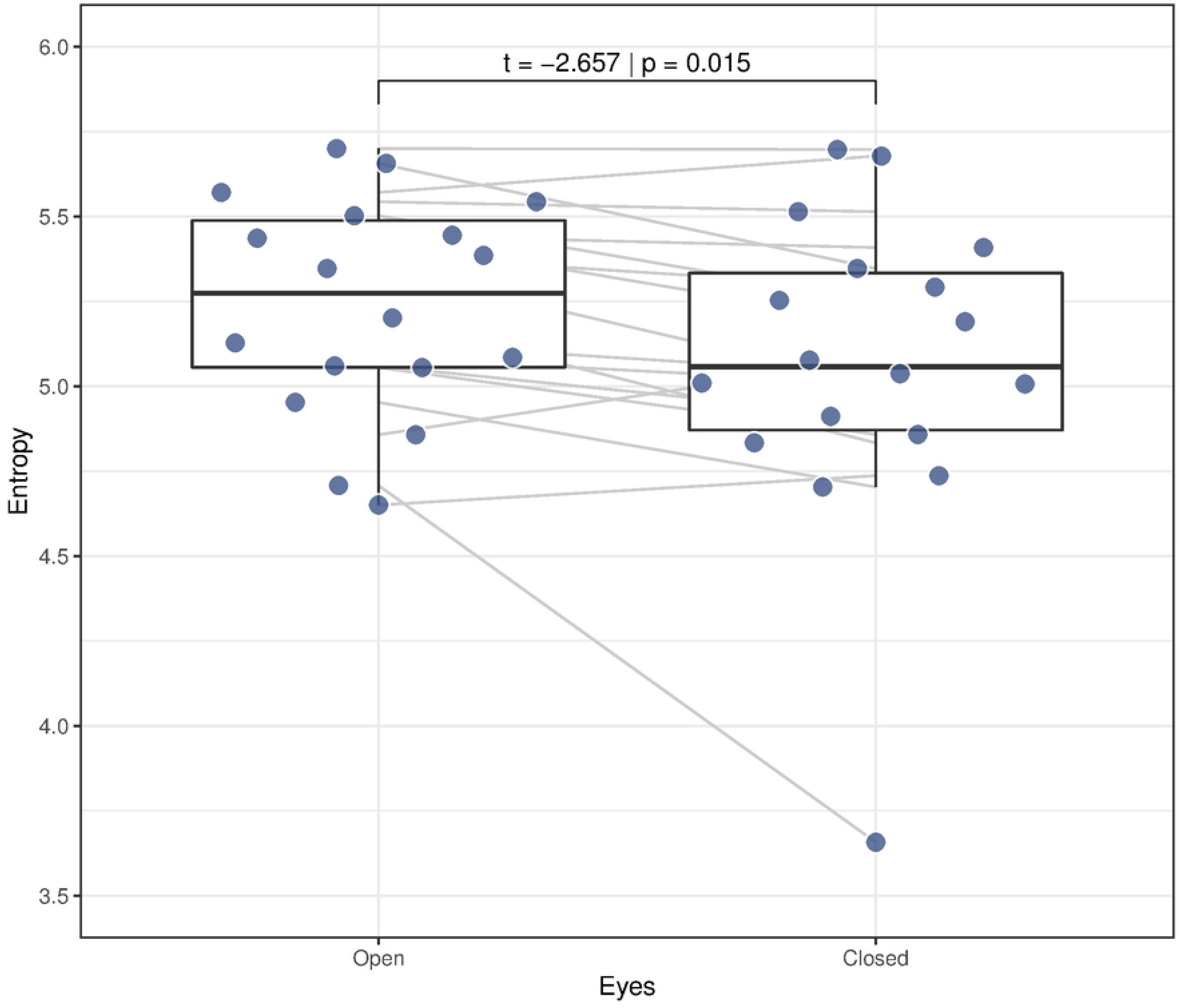
Variation of median entropy with closed eyes. A reduced cortical entropy is associated with closed eyes, the variation is statistically significant as evaluated with the GLM1: *MedianEntropy* ∼*Eyes*(*Open/Closed*) + (1 | *Subject*) and a two-sided 10,000 repetitions permuation test.

### Cycling is associated with reduced cortical entropy

We then analysed the effect of cycling in the electrophysiological signal. The cycling state shows lower entropy when compared with the resting state for left and right central (*t* = ™3.824; *p* = 0.001), left and right frontal (*t* = −3.617; *p* = 0.001 and *t* = −2.164; *p* = 0.042, respectively), left and right occipital (*t* = −3.337; *p* = 0.002 and *t* = −2.901; *p* = 0.006, respectively), and left and right parietal (*t* = −4.754; *p <* 0.001 and *t* = −2.497; *p* = 0.018, respectively) regions, shown in Fig 2A.

**Fig 2.**
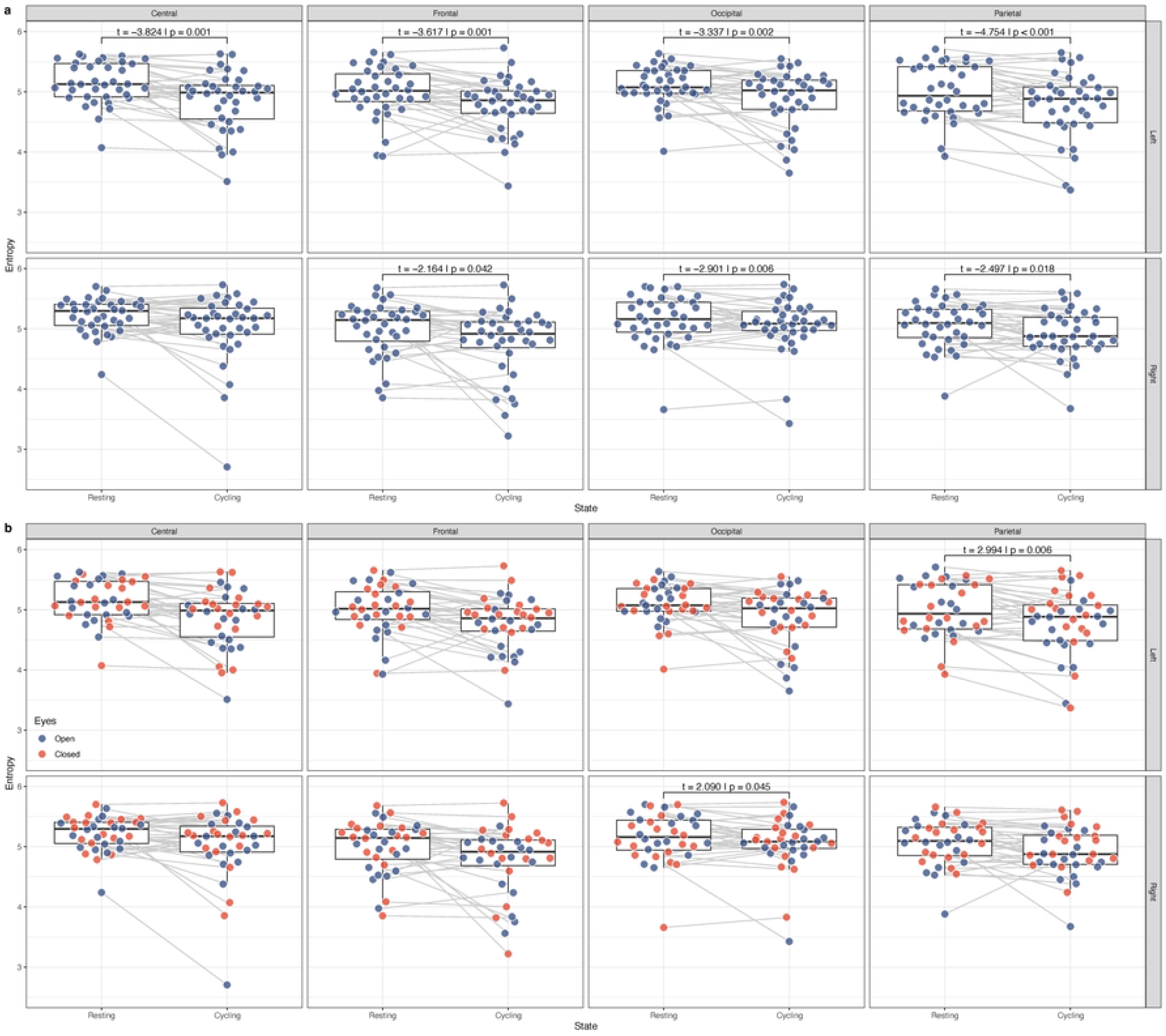
Variation of median entropy with cycling. a) A reduced cortical entropy is associated with cycling. b) Measures for closed and open eyes session. A reduced cortical entropy is also associated with interaction between cycling and having one’s eyes closed. Reported values are statistically significant as evaluated with the GLMM2: *MedianEntropy* ∼ *State*(*Cycling/Rest*) * *Eyes*(*Open/Closed*) + (1| *Subject*) and a two-sided 10,000 repetitions permuation test.

### Cycling with closed eyes and cortical entropy

To evaluate the effect of cycling with closed eyes, we have also analysed the interaction term for our model for closed eyes and cycling. The cycling state with eyes closed shows lower entropy for right occipital (*t* = 2.090; *p* = 0.045) and left parietal (*t* = −2.994; *p* = 0.006) regions, shown in Fig 2B. A topographic plot with the entropy profile for the four different scenarios is shown in Fig 3.

**Fig 3.**
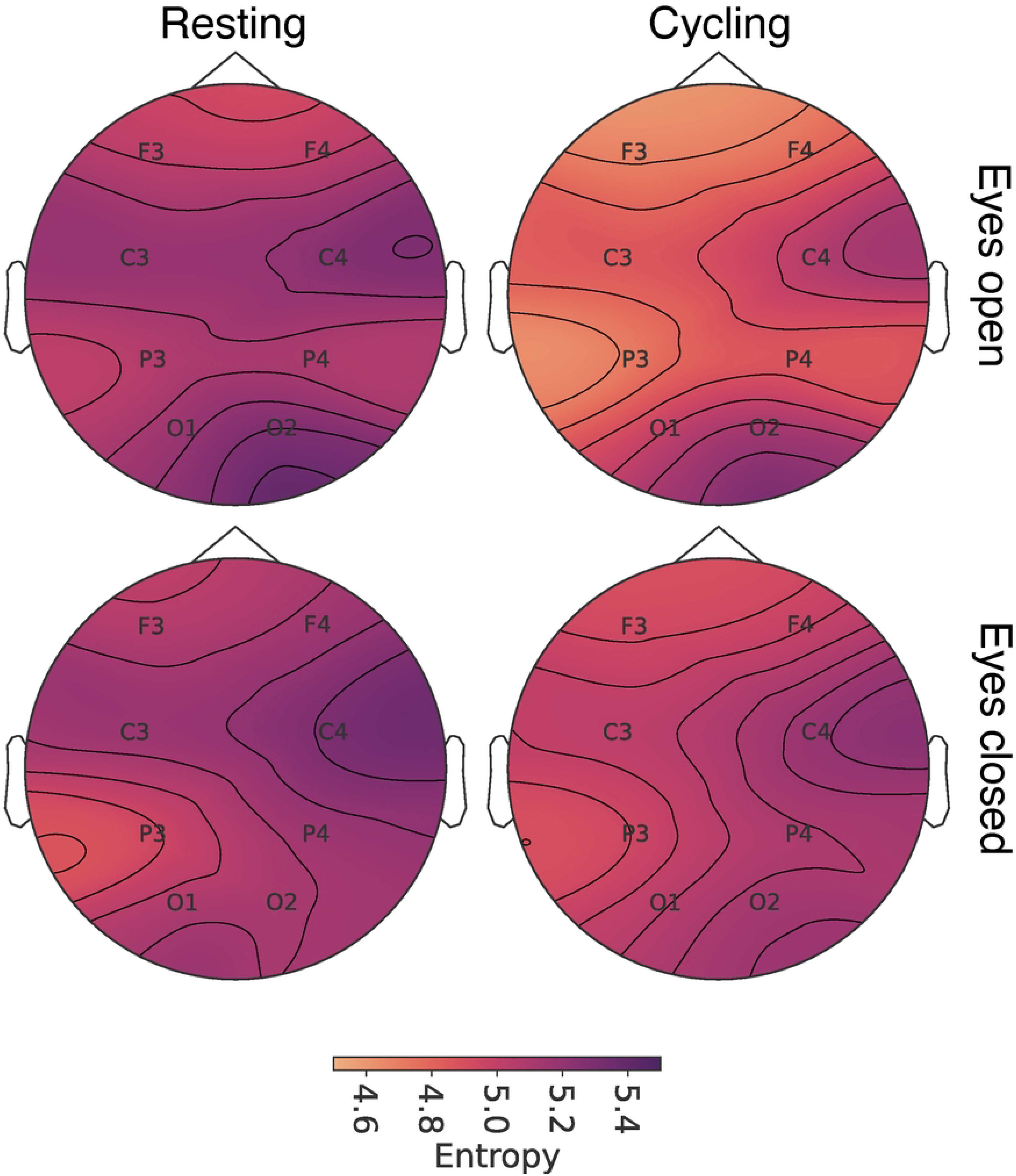
Topographic plots for median entropy on F3-Fz, F4-Fz, C3-Cz, C4-Cz, P3-Pz, P4-Pz, O1-A2 and O2-A1 electrodes. The profiles show a reduction in entropy figures for eyes closed and cycling states - with lowest values registered for cycling with eyes closed.

## Discussion

We applied the recurrence entropy [27, 28] as a measure of complexity to assess changes in cortical electrophysiological activity associated with distinct brain functional states. The methodology for estimating signal complexity via entropy of recurrence was initially validated using the electrophysiological signal of the occipital region and comparing the signal entropy between open-eyes and closed-eyes. The result showed a reduction in entropy in the closed-eyes condition. We know that the open-eyes condition allows retinal light detection which leads to neural activity in the occipital region responsible for processing visual information, that is represented by desynchronised neuronal activity. When we close our eyes and thus prevent light stimuli, an intrinsic neural activity of the thalamus-cortex circuit dominates the signal with a predominant frequency between 8 and 12 Hz, known as the alpha rhythm [32, 33]. Our results suggest that the complexity measurement we propose is capable of distinguishing both: a synchronised pattern with low entropy (closed eyes) and a desynchronised pattern with high entropy (open eyes).

The aim of our study was to evaluate electrophysiological changes between rest and cycling conditions. Using the same recurrence entropy, we showed that the neural complexity is lower during the cycling behavior when compared to the resting state.

This reduction in entropy was observed with greater predominance in the anterior areas of the brain, especially in the frontal area.

We suggest that the observed entropy reduction is associated with an increase in cortical synchronization due to the increase of sensory feedback originating from the lower limbs as a result of flexion and extension repetitive movements that alternately occur in the lower limbs during cycling. Our results are in line with those from Storzer et al. [10] who demonstrated a different pattern in the dynamics of neuronal oscillation in the cortex associated with walking and cycling behaviors. Cycling behavior reduces the neuronal activity at high frequencies, in the beta band range (20 to 30 Hz) during the movement, followed by a rebound of this beta activity when exercise terminates [10]. During cycling, pedals are held together, and thus both legs are alternately moving. To understand these neuronal dynamics during walking behavior, the authors suggest that during walking, unlike cycling, each leg is independent with alternating stance and swing phases; that is, the movement is divided into short, distinct independent motions. In this way, decreased beta potency has been linked to an active neural state in the sensorimotor cortex that is associated with an increase in cortical excitability [34].

Furthermore, it has been shown that the beta power remains suppressed during continuous movements [35], but it is maintained in isometric and sustained movements [36, 37]. As the cycling is continuous, this movement may cause a stronger decrease in beta power, and it would explain the entropy reduction found in our study.

Spectral analysis is widely used by neuroscientists and indeed the identification of brain waves is built upon frequency bands [38]. Nevertheless, this technique has some important drawbacks, for example the dependence on noise that pollutes the frequency spectrum, often making it difficult to identify the dominant frequencies [39]. In addition, the EEG signal is non-linear and non-stationary with a high degree of complexity [40], and therefore the use of the Fourier transform is not entirely appropriate [16]. Furthermore, the spectral analysis is dependent on some arbitrary factors of choice: the size and shape of the window to analyse the signal and the type of base used. We point out that the basis of sines and cosines associated with the Fourier transform is the most common basis, but the Wavelet transforms open the way to many alternative bases [41]. In recent years, with the availability of mathematical tools based on complexity theories [22, 23], we have observed the use of entropy-based approaches as a strategy for nonlinear EEG analysis to provide independent and complementary measurements to conventional EEG spectral analysis, and with this, it has been possible to characterise these entropy measures with discrete changes in behavioral states [15, 42]. In this context, the recurrence entropy is a tool that directly uses the time series, without the need for preprocessing involving spectral analysis, to estimate the complexity of an electrophysiological signal.

## Conclusion

Our results concerning the decrease in neuronal complexity highlight the potential of recurrence entropy as a tool to study the neural mechanisms involved in Parkinson’s disease [43, 44]. In particular, this tool could be useful to reveal the ability to ride a bicycle that even patients with severe gait freezing present [13, 45]. Parkinson’s disease is commonly characterized by an increased beta band activity in the neural circuit of the basal ganglia [46]. Freezing gait in Parkinson’s patients has been explained by impaired temporal and spatial gait control [47], with a decrease in neural activity in the supplementary motor area [48] and also by an increase in high beta activity during the resting state in the subthalamic nucleus [49]. Thus, we conjecture that cycling is not impaired in Parkinson’s disease due to the continuity of sensory feedback caused by repetitive movements that promote a reduction in entropy, a decrease in neuronal complexity associated with less sensorimotor processing. In future work we intend to use this technique to assess brain complexity in different cycling conditions, for instance, on a horizontal and vertical stationary bicycle or cycling in a virtual reality environment. In addition, we have the perspective to explore the brain complexity of subjects who know, or do not, how to ride a bicycle when driving a stationary bicycle.

## Acknowledgments

The authors thank the Brazilian Council for Scientific and Technological Development (CNPq) for its continued support. Grants Nos. 309440/2022-0, 308441/2021-4, 305189/2022-0, grant 307907/2019-8.

